# XPO1 inhibition modulates the Wnt/β-catenin signaling pathway to reduce colorectal cancer tumorigenesis

**DOI:** 10.1101/2024.10.31.621312

**Authors:** Andrew E. Evans, Sahida Afroz, Alexa Magstadt, Dan A. Dixon

## Abstract

Colorectal Cancer (CRC) is the second leading cause of cancer-related death in the U.S. and high-risk individuals face a notably higher likelihood of developing CRC based on their genetic background. Hence, there is a compelling need for innovative chemopreventive treatments aimed at minimizing CRC tumorigenesis. Exportin 1 (XPO1; also referred to as CRM1) plays a pivotal role in transporting proteins from the nucleus to the cytoplasm. Various cancers overexpress XPO1, including CRC, and Selective Inhibitors of Nuclear Export (SINE) compounds, such as Eltanexor (KPT-8602), have been developed to target XPO1. Eltanexor demonstrates fewer adverse effects than its precursors and is currently under evaluation in Phase I/II clinical trials. This research evaluates Eltanexor as a chemopreventive agent for CRC. Our findings indicate Eltanexor treatment inhibits expression of the common chemoprevention target in CRC, cyclooxygenase-2 (COX-2). This occurs by Eltanexor-dependent reduction of Wnt/β-catenin signaling. Furthermore, XPO1 inhibition leads to forkhead transcription factor O subfamily member 3a (FoxO3a) nuclear retention, which can modulate β-catenin/TCF transcriptional activity. *In vivo* oral treatment of Eltanexor to Apc*^min/+^ mice (a mouse model for Familial Adenomatosis Polyposis) was well-tolerated and* reduced tumor burden by approximately 3-fold, along with decreased tumor size. Drug sensitivity assays using organoids from Apc*^min/+^ mice* tumors show increased sensitivity to Eltanexor compared to wild-type organoids. Collectively, these findings highlight XPO1 as a potent target for CRC chemoprevention.

**SIGNIFICANCE:** In this study, we show the XPO1 inhibitor, Eltanexor, reduces COX-2 by modulating the Wnt/β-catenin signaling pathway and acts as an effective chemopreventive agent in the Familial Adenomatous Polyposis (FAP) mouse model, Apc*^min/+^* mice.

## INTRODUCTION

In the United States, CRC is the second leading cause of cancer deaths in the U.S., affecting both men and women. The American Cancer Society predicts that 53,010 people will die from CRC in the year 2024[1]. These statistics highlight the clear need for novel treatment approaches to combat CRC. This need is particularly true due to the prevalence of CRC, which is expected to rise in young people. People born in the year 1990 have double the risk of developing colon cancer when compared to someone born in the year 1950[2]. In addition to younger people being more likely to develop CRC in their lifetime, they are also more likely to experience an early onset of CRC. Since 1994, the incidence of early-onset CRC (individuals younger than 50 years old) has been increasing by about 2% each year[3] due to numerous risk factors.

In addition to the sporadic development of CRC, individuals with conditions such as Familial Adenomatous Polyposis (FAP) are predisposed to CRC development due to inherited germline mutations[4]. For individuals diagnosed with FAP, clinical providers recommend that they begin annual colonoscopies at the age of 10-12 years old. For people with FAP, the risk of CRC is 100%, thus, forcing many of these patients to receive a colectomy to prevent CRC[5]. Given the heightened CRC risk faced by these individuals, we must develop chemopreventive agents to reduce their risk of CRC and improve their quality of life.

The nuclear export protein, Exportin 1 (XPO1; also known as CRM1), is crucial in transporting proteins with a leucine-rich Nuclear Export Signal (NES) from the nucleus to the cytoplasm[6]. XPO1 is overexpressed in multiple different cancer types, including colon cancer[7]. The upregulation in XPO1 expression can result in the excessive removal of over 1000 different proteins from the cell’s nucleus. Included in those 1000+ proteins are proteins known to be associated with the development of CRC[8,9]. To target XPO1, a novel class of drugs known as Selective Inhibitors of Nuclear Export (SINE) compounds have been developed[10]. A second generation of SINE compound, Eltanexor (KPT-8602), is currently in Phase I/II clinical trials for multiple cancer types while demonstrating fewer side effects than precursor SINE compounds[11] (ClinicalTrials.gov <u>NCT02649790</u>). XPO1 inhibition impairs numerous hallmarks of cancer, including the ability to induce DNA damage and apoptosis; while reducing inflammation, cell proliferation, and angiogenesis [12–16].

Given that XPO1 is overexpressed in CRC and its inhibition reduces numerous hallmarks of cancer, we hypothesize that Eltanexor will act as an effective chemopreventive agent.

This report shows that Eltanexor effectively limits cell viability in CRC cells, reduces the chemoprevention target Cyclooxygenase-2 (COX-2), and impairs the transcriptional activity of a paramount signaling pathway in CRC, Wnt/β-catenin signaling. In-vivo, we show that Eltanexor effectively reduces tumor burden in a FAP mouse model, Apc*^min/+^* mice.

## MATERIALS AND METHODS

### RNA Analysis

Colon cancer cDNA Array was purchased from Origene (Cat#: HCRT104). For normal colon tissue used in the CRC cell array, normal tissue cDNA was obtained from Origene and normalized to GAPDH for analysis. For all other RNA analysis, samples were normalized to actin. Total RNA from cell culture was extracted using the Qiagen RNeasy Mini Kit (Cat# 74104). cDNA synthesized using 1 μg of total RNA in combination with olgio(dT) and Improm-II reverse transcriptase (Promega; Cat# A3801). qPCR was performed as previously described[17] with SYBR green PCR master mix (Applied Biosystems; Cat# A25778) on 7300 PCR assay system (Applied Biosystems). Fold in mRNA expression levels was analyzed as previously described[18].

**Table 1.**
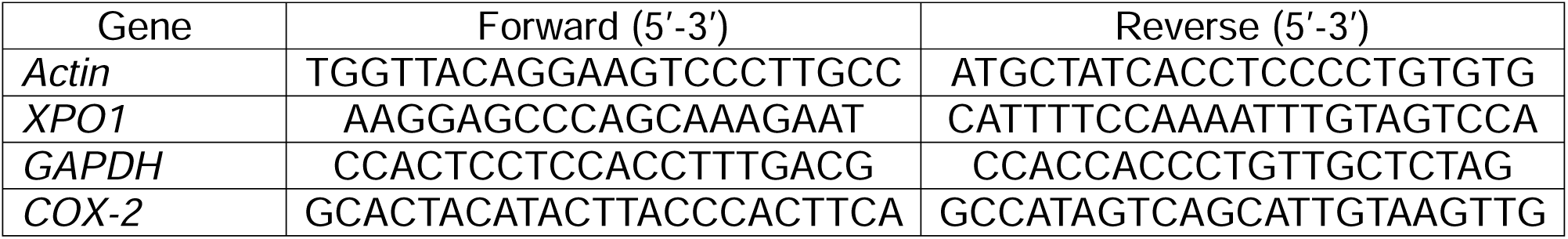
qPCR primer sequences.

### Cell Culture

Colorectal cancer cells (DLD1, HT-29, HCT116, MOSER, SW480, Caco2, and RKO) were purchased from American Type Culture Collection (ATCC). HCA-7 cells were provided by S. Kirkland (Imperial College, London, United Kingdom). DLD1, HT-29, HCT116, HCA-7, MOSER, SW480, Caco2 were cultured in Dulbecco’s Modified Eagle’s Medium (Corning-USA; Cat# 10-013-CV) and RKO cells were culture in Minimum Essential Medium (Corning-USA; Cat# 10-009-CV). All media was supplemented with 10% Fetal Bovine Serum (R&D systems; Cat# S11150) and 100 units per ml Penicillin-Streptomycin (Corning; Cat #30-002-CI). Cell lines were not mycoplasma tested. All cell lines were between passage 2-25. The cells lines DLD1, HCT116, SW480, Caco2, and MOSER are derived from males. The cell lines HCA-7, HT-29, and RKO are derived from females. Phase-contrast images were taken using an EVOS Fl Auto Imaging System. Eltanexor (APExBIO; Cat # B8335) was dissolved in DMSO (Sigma; Cat# 34869) and diluted in complete media.

### Cell Viability Assays

CCK8 assays was used to analyze cell viability. 1000 cells were plated into each well of a 96 well plate. Twenty-four hours later, cells were treated with varying concentrations of Eltanexor (APExBIO). After 72 hours, 10 μL of CCK8 (GLPBIO; Cat # GK10001) was added to each well and the plate incubated at 37 °C for 1 hour before reading each well absorbance at 450 nm on a BioTek ELX800 Microplate reader.

For the colony formation assay, cells were seeded at 500 cells/ well in 6-well plates and treated with 800 nM Eltanexor (APExBIO), 200 nM Eltanexor, or a DMSO-vehicle (Sigma) control. After 72 hours, the drug treatment medium was replaced with complete DMEM. DMEM was changed every three days for the remainder of the experiment. 11 days after initial treatment, cells were washed with PBS, incubated for 20 min in 100% methanol, and stained for 40 min in .1% crystal violet solution. Colonies were analyzed for percentage of total plate coverage utilizing the ColonyArea ImageJ plugin: https://imagej.net/plugins/colonyarea

### Fluorescence Microscopy

Cells were grown on coverslips and treated with varying dosages of Eltanexor for 24 or 48 hours. Following the completion of treatment, cells were washed once with PBS and fixed with 4% formaldehyde (Fisher; Cat# F79-500) for 15 minutes. Cells were then washed with PBS and blocked with PBS + 5% normal goat serum (Cell Signaling; Cat# 5425S) + 0.3% Triton X-100 (Sigma; Cat# T8787) for 1 hour. The blocking buffer was removed, and cells were incubated with either anti-XPO1 (1/400; Cell Signaling Cat # 46249S), anti-FoxO3a (1/400; Cell Signaling Cat # 12829S), or anti-β-catenin (1/200, Cell Signaling, Cat # 8480S) diluted in PBS + 1% Bovine Serum Albumin (Fisher) + 0.3% Triton X-100 overnight at 4°C. Cells were then rinsed 3x with PBS. Secondary antibody was applied to the cells for 1 hour at room temperature using Alexa Fluor 488 goat anti-rabbit (1/500; Invitrogen, Cat# A11008) diluted in the antibody dilutant. Cells were then counterstained with DAPI (Santa Cruz; Cat# sc-24941) and imaged using a EVOS Fl Auto Imaging System at 60x magnification.

For immunofluorescence on tissue, the tissue was formalin-fixed paraffin embedded (FFPE). 5 μm slides were cut and rehydrated by washing the samples 5 minutes/ wash with xylene 2x, 100% ethanol, 95% ethanol, 80% ethanol, 50% ethanol, diH_2_O, and PBS. Antigen retrieval was performed microwaving slides in antigen citrate buffer pH 6.0 (Sigma; Cat# C9999) for 1 minute at high-power and 5 minutes a low power. The samples were then placed in a steamer for 30 minutes, and then placed back in antigen citrate buffer pH 6.0 for 20 minutes. The slides were then washed for 5 minutes in PBS. Blocking buffer made of TBST + 5% NGS (Cell Signaling) was applied to the slides for 2 hours. Slides were then washed with PBS for 3 minutes and primary antibody diluted in blocking buffer was applied to the slides overnight at 4°C. Antibodies used include: anti-XPO1(1/200; Cell Signaling, Cat# 46249S), anti-COX-2 (1/300; Cell Signaling, Cat# 12282S), anti-Ki67 (1/200; Cell Signaling, Cat# 12202S). Slides were then washed with PBS 3x, and Alexa Fluor 488 goat anti-rabbit (1/500; Invitrogen, Cat# A11008) diluted in blocking buffer was applied for 2 hours. Slides were washed 3x with PBS and then incubated with Hoechst 33342 (1/1000; Invitrogen, Cat# H3570) diluted in PBS for 5 min. The slides were washed 3x in PBS and incubated in 0.1% Sudan Black (Sigma; Cat# MKBB2665) diluted in 70% ethanol for 20 minutes. Slides were washed 3x with PBS and mounted using Prolong Gold Antifade (Invitrogen; Cat# P36930). Slides were imaged using EVOS Fl Auto Imaging System at 20x magnification.

### Tumor Growth Xenograft

HCT116 cells were thawed and passaged twice before the day of injection. On the day of injection, cells were trypsinized, counted, and 20 million cells/ mL were resuspended in PBS. The cells were then injected into the dorsal flanks of athymic nude mice (The Jackson Laboratory). Three days post-injection, the mice were randomly separated into either vehicle-treated or Eltanexor-treated groups. The vehicle was comprised of 0.5% methylcellulose (Sigma; Cat# M0262) and 1% Tween-80 (Sigma; Cat# P4780). Eltanexor (APExBIO) was dissolved in the vehicle. Mice were treated with the vehicle or 10mg/kg of Eltanexor 3x/ week by oral gavage. Tumor volume was monitored 3x/ week by caliber. Tumor volume was calculated using the formula tumor length x (width)^2^/2. Once the tumors reached 1200-1500 mm^3^ in size, the mice were sacrificed, the tumors were excised, and photographs were taken. The tumors were formalin-fixed paraffin-embedded for immunohistochemistry analysis.

### Western Blot Analysis

Western blots performed as described[19]. The primary antibodies used includes: anti-XPO1 (1/1000; Cell Signaling, Cat# 46249S), anti-FoxO3a (1/1000; Cell Signaling, Cat# 12829S), anti-c-myc (1/1000; Cell Signaling, Cat# 5605S), anti-COX-2 (1/1000; Cell Signaling, Cat # 12282S), anti-Actin (1/5000; MP Biomedical, Cat # 691002), anti-β-catenin (1/1000, Cell Signaling, Cat # 8480S).

### DNA and siRNA transfections

Evaluation of the COX-2 promoter was done through luciferase assays utilizing pGL3-COX2(−1840), pGL2-COX2(−1800), pGL2-COX2(−1400), pGL2-COX2(−800), pGL2-COX2(−400), pGL2-COX2(−300). Our generation of these plasmid constructs is previously described. Furthermore, TOPflash luciferase reporter construct was used evaluate Wnt/ β-catenin signaling as previously described. The plasmid constructs were transfected using Lipofectamine LTX and PLUS (Invitrogen; 15338100). One day post transfection, cells were treated with varying doses of Eltanexor (APExBIO) for 24 or 48 hours. The cells were then lysed and assayed as previously described[18].

For FoxO3a siRNA KD, FoxO3a siRNA (Horizon; Cat# L-003007-00-0005) and siRNA Control (Ambion; Cat# AM4635) were transfected into cells using siQuest (Mirus; Cat# 2114). The cells were transfected with the siRNA for 24 hours, were then left to recover for 24 hours, transfected again for 24 hours, and then assayed 48 hours post-second transfection.

### Organoid Growth and Viability Assays

To establish mouse organoid cultures, 15-week-old Apc*^min/+^*or 15-week-old wild-type (WT) C57BL/6 mice were euthanized. The most proximal 20cm of the small intestines of the mice were removed, washed with PBS, and longitudinally splayed open. For the Apc*^min/+^* mice, between 20-30 tumors were excised. For the WT mice, the small intestine was chopped into 2cm segments using a razor in PBS. Tissue was then washed 3x with D-PBS (Corning; Cat# 21-0310-CV), 3x with D-PBS + 2% Penicillin-Streptomycin (Sigma), and then rocked for 90 minutes at 4°C with dissociation buffer containing 3mM EDTA (Invitrogen; Cat# AM9260G) and 0.5mM DTT (Sigma; Cat# 10197777001) in D-PBS. Post dissociation, dissociation buffer was removed, and 10 mL of D-PBS was added. Tissue was pipetted 10 times; tissue was allowed to gravity settle for 30 seconds, then supernatant was removed and filtered through a 70u cell strainer (PR1MA; Cat# 70ICS). This process was repeated 3 more times to create a total of 4 fractions. The fractions are then spun at 300xg for 5 minutes (Eppendorf 5804 R) at 4°C. Pellets were resuspended in 10mL of cold PBS+ 0.1% (w/v) BSA (Fisher; Cat# 9048-46-8). The fractions are then spun at 200xg for 3 minutes at 4°C and pellets are resuspended in 5mL DMEM/F12 (Cytiva; Cat# SH30023.01). For WT the number of isolated crypts were counted in a 10 μL sample to calculate number of crypts/ mL. Furthermore, 4000 crypts were centrifuged at 200xg for 5 min at 4°C. Pellets were resuspended in 150 μLof Mouse IntestiCult Organoid Growth Medium (STEM; Cat #06005). Then, 150 μL of Matrigel (Corning GFR, Phenol-red free, LDEV Free; Cat #CB-40230C) was added to each tube, and 50μLof crypt solution was plated into each well of a pre-warmed 24 well tissue culture non-treated plate (Falcon; Cat# 351147). After 15min, 750 μL of Intesticult Media was added to each well. Media was then swapped every 2-3 days and organoids were stored in an incubator kept at 37°C and 5% CO_2_.

On day 5, both WT organoids and Apc*^min/+^* tumor organoids were treated with varying doses of Eltanexor (APExBIO) or DMSO (Sigma). The organoids were treated for a duration of 72 hours. Every 24 hours, photos of the organoids at different treatment concentrations were captured using a EVOS Fl Auto Imaging System microscope at 4x magnification. Organoid size was quantified using ImageJ version 1.53k (NIH). After 72 hours, organoid media was removed and replaced with PBS containing 10 μg/mL of Propidium Iodide (Sigma; Cat# S7109) and Hoechst 33342 (Sigma). The organoids with PBS solution incubated at 37°C and 5% CO_2_ for 30 minutes and images of organoids on blue and red fluorescent channels using a EVOS Fl Auto Imaging System microscope at 4x. The number of PI-positive organoids were then scored.

### Animals

All procedures were reviewed and approved by the University of Kansas Institutional Animal Care and Use Committee. The Apc*^min/+^* and wild-type mice utilized were in the C57BL/6J background and originally purchased from Jackson Laboratory. The colonies were maintained in a pathogen-free environment at the University of Kansas Animal Care Facility.

In the Eltanexor treatment study, the vehicle used was 0.5% methylcellulose (Sigma) and 1% Tween-80 (Sigma). 7 Apc*^min/+^* mice received vehicle treatment and 7 Apc*^min/+^* mice received 10mg/kg Eltanexor (APExBIO) dissolved in vehicle and administered by oral gavage. The dosage of 10mg/kg was determined based on previous studies[7,20]. The sex, weight, and age of the mice prior to the start of the study was balanced. Eltanexor or vehicle administration occurred 3x/ week and began when the mice were between 4-5 weeks old and lasted for a duration of 6 weeks. On each day the mice received treatment, weights were recorded. Upon conclusion of the study, the mice were euthanized with isoflurane and the intestinal tract was removed washed with PBS, longitudinally splayed, and formalin-fixed in 10% formalin (Fisher; Cat# SF100) for 24 hours and then stored in 70% ethanol. Stored tissue was given an identification code to conceal the treatment condition. Furthermore, spleens from the mice were removed and weighed.

After tissue was stored in 70% ethanol, small intestine and colon tissue was examined under a dissecting scope (Olympus SZX12). Total number of polyps and polyp sizes were determined by two individual investigators who were blinded to the treatment conditions. After counting and sizing tumors, the intestinal tissue was formalin-fixed paraffin-embedded.

### Immunohistochemistry

FFPE tissue was cut into 5 µm sections. Sections were then rehydrated through a series of xylene 2x, 100% ethanol, 95% ethanol, 80% ethanol, 50% ethanol, diH_2_O, and PBS washes at 5 minutes/ wash. Antigen retrieval (Sigma) was performed on the slides, as described under the immunofluorescence section. Slides were then washed with PBS for 5 min. Then, the slides were exposed to H_2_O_2_ for 10 minutes. The slides were then washed with PBS and blocked for 2 hours using TBST + 5% NGS (Cell Signaling). Slides were then exposed to anti-XPO1 (1/200; Cell Signaling Cat # 46249S) or Ki-67 (1/200; Cell Signaling, Cat# 12202S) overnight at 4°C. The next day, slides were washed 2x with PBS and anti-rabbit secondary antibody (SignalStain Boost IHC Detection Reagent HRP Rabbit; Cell Signaling, Cat# 8114) was applied for 2 hours. The samples were then washed 2x with PBS for 5 min/ wash and for 3 minutes with dH_2_O. DAB solution (Cell Signaling; Cat# 8059S) was applied to visualize the peroxidase reaction. Slides being compared were exposed to DAB chromogen for equal durations of time. Slides were then counterstained with Hematoxylin 560MX (Leica; Cat# 3801576) and mounted using cytoseal XLT (Thermo Scientific; Cat# 8312-4)

### Hematoxylin and Eosin Staining

FFPE tissue was sectioned into 5 µm sections. Slides were rehydrated through a series of washes in xylene and ethanol. The nuclei were stained with Hematoxylin 560MX (Leica). Bluing (Leica; Cat# 3802915) and Differentiating buffers (Leica; Cat# 3803595) was applied to slides. The cytoplasm was then counterstained with Eosin (Leica; Cat# 3801615). Slides were mounted using cytoseal XLT (Fisher) and imaged with an Nikon Eclipse TE2000-U microscope.

### Statistical Analysis

All graphs and statistical analysis performed in this paper was done using Graphpad Prism v10.1.1. When applicable, unpaired students t-test was performed with values ≤

### 0.05 considered statistically significant

### Data Availability

The data generated in this study are available upon request from the corresponding author.

## RESULTS

### Targeting the overexpression of XPO1 in CRC

To assess XPO1’s expression in CRC tumor tissue compared to normal colon tissue, we analyzed The Cancer Genome Atlas Program (TCGA) data through the Gene Expression Profiling Interactive Analysis (GEPIA). The GEPIA data shows that XPO1’s gene expression is elevated in colon adenocarcinoma (COAD) tissue compared to normal colon tissue (**Figure 1A**). Furthermore, we examined XPO1’s expression amongst stage 1-4 CRC, XPO1 is consistently expressed (**Figure 1B and Figure S1 A**). The consistent overexpression of XPO1 further suggests that the overexpression is an early event in CRC. When we examine XPO1 gene expression in multiple CRC cell lines, the protein is consistently overexpressed (**Figure 1C**). This data concurs with previous reports showing XPO1 protein is overexpressed in various CRC cell lines [7]. When we examined how XPO1 expression affects overall CRC survival, we observed increases in XPO1 expression trends towards poorer survival, particularly in microsatellite-stable CRC (**Figure 1S B**). This data, taken together, further suggests the possibility of XPO1 being a viable CRC chemoprevention target. The XPO1 inhibitor and second-generation SINE compound, Eltanexor, inhibits XPO1 by binding to XPO1’s cargo binding pocket, thus blocking XPO1’s ability to interact with cargo proteins (**Figure 1D**).

**Figure 1.**
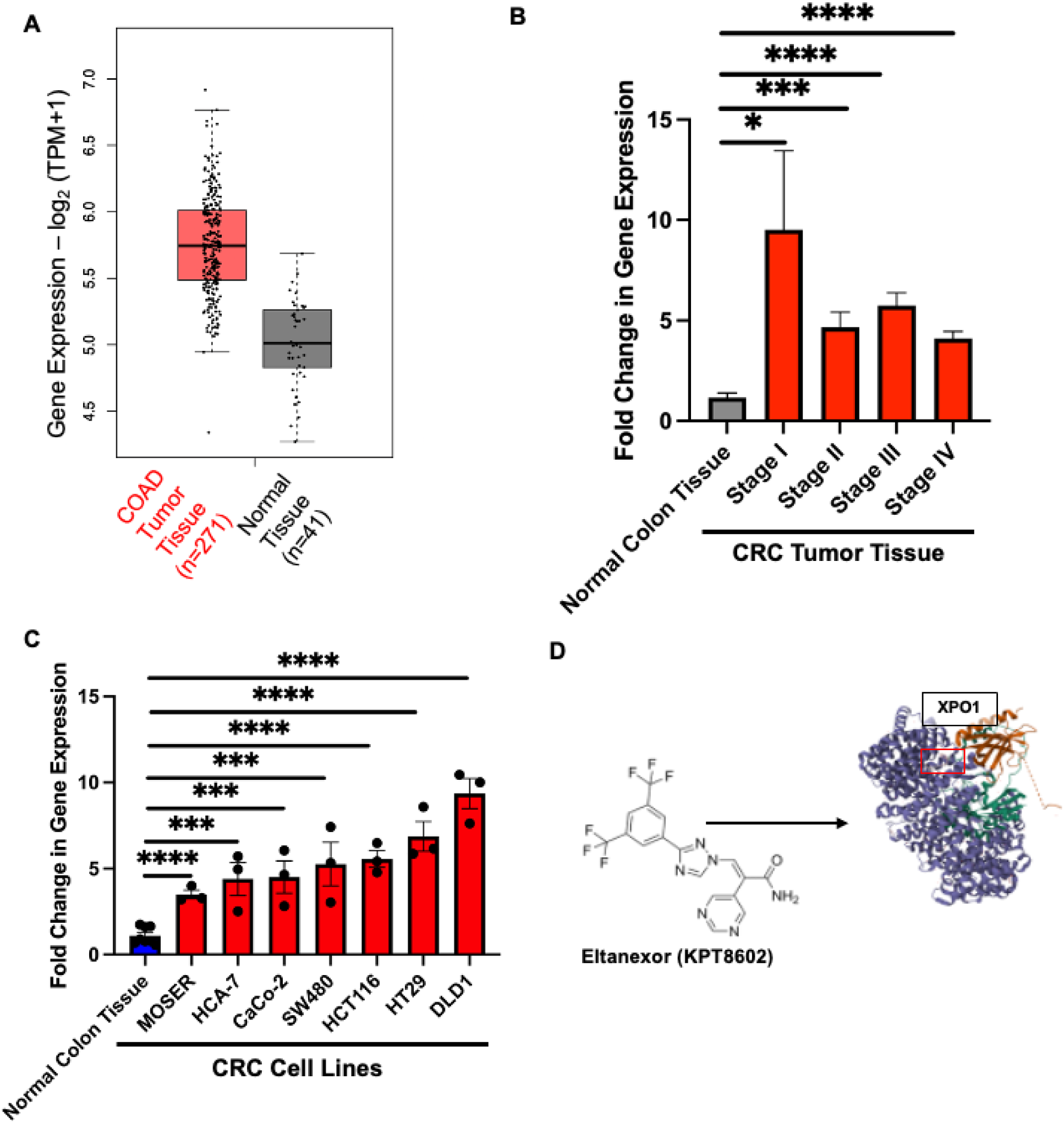
XPO1 is overexpressed in CRC. **(A)** Boxplot derived from GEPIA showing colon adenocarcinoma (COAD) tissue compared to normal colon tissue. (TCGA data derived from online website GEPIA: gepia.cancer-pku.cnn/detail.php?gene=XPO1) **(B)** XPO1 mRNA expression by qPCR in major CRC tumor stages normalized to normal colon tissue. Actin was used as the loading control, and values represent mean fold change ± SEM. **(C)** XPO1 mRNA expression by qPCR in multiple CRC cell lines compared to normal colon tissue. GAPDH was used as the loading control, and values represent mean fold change ± SEM. **(D)** Eltanexor structure and XPO1 structure; highlighting Eltanexor’s protein docking region in XPO1’s protein binding groove. (PDB: 6XJT) *(*, p* ≤ *0.05; **, p* ≤ *0.01; ***, p* ≤ *0.001, ; ****, p* ≤ *0.0001)*

### Eltanexor treatment reduces CRC cell viability and in vivo tumorigenesis

To assess the effectiveness of Eltanexor in CRC, a cell viability assay was performed. This assay demonstrated that Eltanexor exhibits an IC_50_ within the nanomolar range in numerous CRC cell lines with a diverse range of mutational backgrounds (**Figure 2 A and B**). A colony formation assay indicated that Eltanexor reduces CRC cells’ clonogenic ability (**Figure S2 A and B**). To further test if Eltanexor will impair tumorigenesis in an in vivo model, HCT116 cells were injected into the dorsal flanks of athymic nude mice. The Eltanexor-treated mice were given 10mg/kg Eltanexor 3x/week by oral gavage, while vehicle-treated mice received 0.5% Methylcellulose + 1% Tween-80 3x/week by oral gavage. Mice treated with Eltanexor exhibited significantly reduced tumor volume compared to the vehicle-treated mice (**Figure 2 C and D, S3 A**). Eltanexor treatment in the xenograft mice proved to be well-tolerated and reduced the common proliferation marker Ki67 (**Figure S3 A and B**). Furthermore, XPO1 protein expression was examined in CRC cells treated with Eltanexor, revealing a dose-dependent reduction with drug treatment (**Figure 2 E and D, S2 C and D**). XPO1 protein expression was also reduced in vivo in the drug-treated xenograft tumors (**Figure 2 G and H, S3 C**).

**Figure 2.**
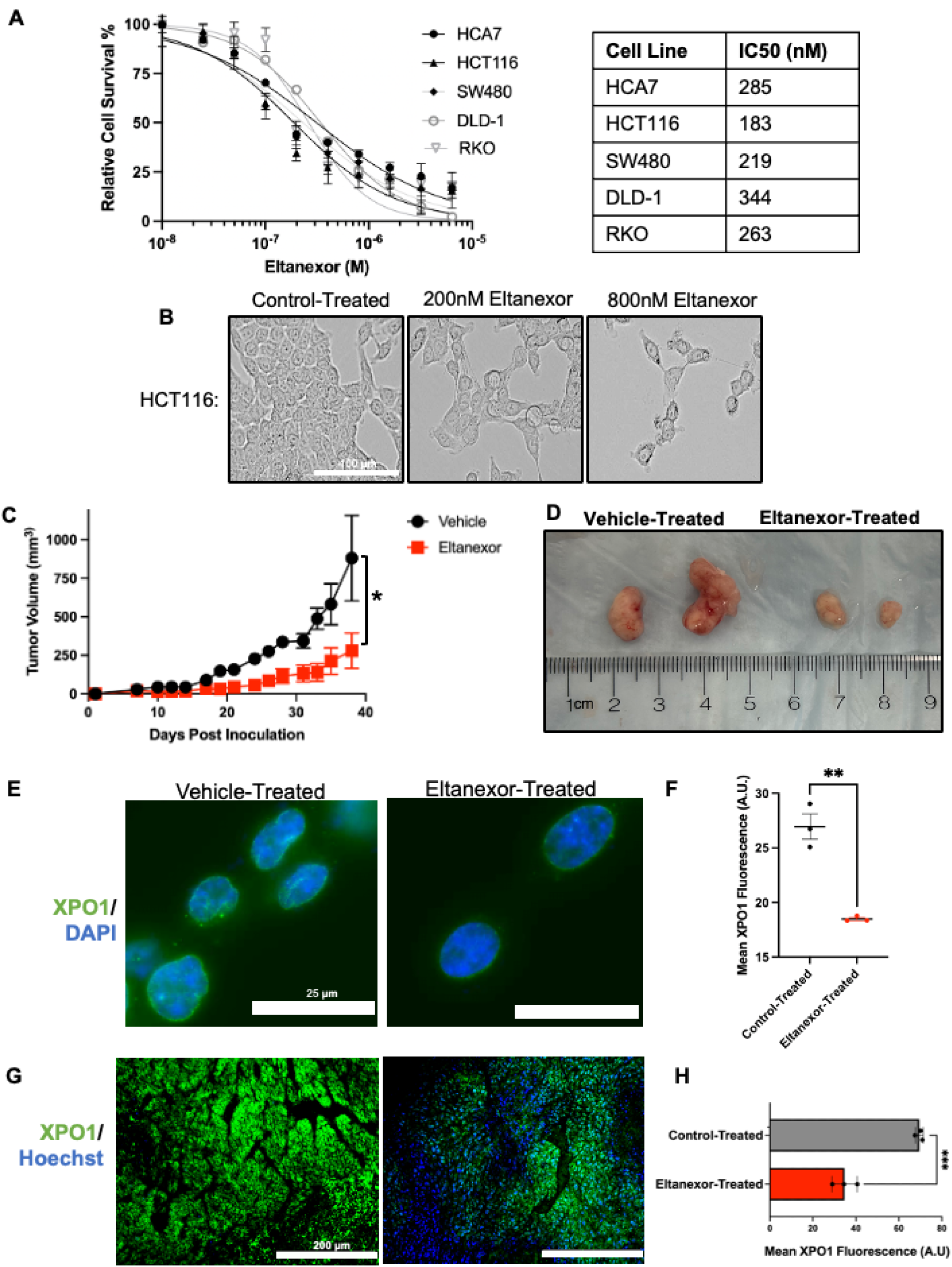
Eltanexor reduces XPO1 protein while reducing cell viability. **(A)** Human colorectal cancer cell lines HCA7, HCT116, SW480, DLD-1, and RKO with varying concentrations of Eltanexor. The cell survival was measured with CCK8 after incubation for 72 hours with Eltanexor. The data represents the mean 7 individual experiments ± SEM. The chart shows determined IC50 values. **(B)** Phase-contrast images showing HCT116 cells treated with varying dosages of Eltanexor for 48 hours. **(C, D)** HCT116 WT cells were injected into the subcutaneous dorsal flanks of athymic nude mice. Measurements were taken 3x a week. Values graphed represent mean tumor volume ± SEM. Mice were either treated with Eltanexor or the vehicle 3x/ week. Representative images of tumors extracted from the vehicle-treated and Eltanexor-treated mice (n= 4 tumors for vehicle-treated/ n=6 tumors for Eltanexor-treated) **(E)** Fluorescent microscopy image of HCT116 cells treated with control or Eltanexor for 48 hours. Green represents XPO1, and DAPI was used to stain the nucleus. The scale bars represent 25μm. **(F)** Comparison of XPO1 fluorescent signal between control-treated and Eltanexor-treated cells. The values graphed are the mean fluorescence values of 3 independent experiments ± SEM. At least 40 cells were quantified for each experiment. **(G)** Immunofluorescent detection of XPO1 in the vehicle and Eltanexor-treated tumors. Green represents XPO1 and Hoechst was used to stain the nucleus. Representative tissue sections were used and imaged at 20x magnification. The scale bars represent 200μm **(H)** Comparison of XPO1 fluorescent signal between vehicle-treated and Eltanexor-treated tumors. The values graphed are the mean fluorescence value of 3 xenograft tumors ± SEM. *(*, p* ≤ *0.05; **, p* ≤ *0.01; ***, p* ≤ *0.001)*

### COX-2 is reduced with Eltanexor treatment

To thoroughly investigate Eltanexor as a chemopreventive agent, we examined whether Eltanexor treatment altered cyclooxygenase-2 (COX-2) expression. COX-2 inhibition through inhibitors such as nonsteroidal anti-inflammatory drugs (NSAIDS) are frequent chemoprevention treatment options for individuals at high risk for CRC[21]. Furthermore, we have shown that COX-2 is overexpressed in CRC[22]. Interestingly, when we treated HCA-7 cells with 200nM Eltanexor for 48 hours, we observe a reduction in COX-2 protein and gene expression (**Figure 3 A and B**).

**Figure 3.**
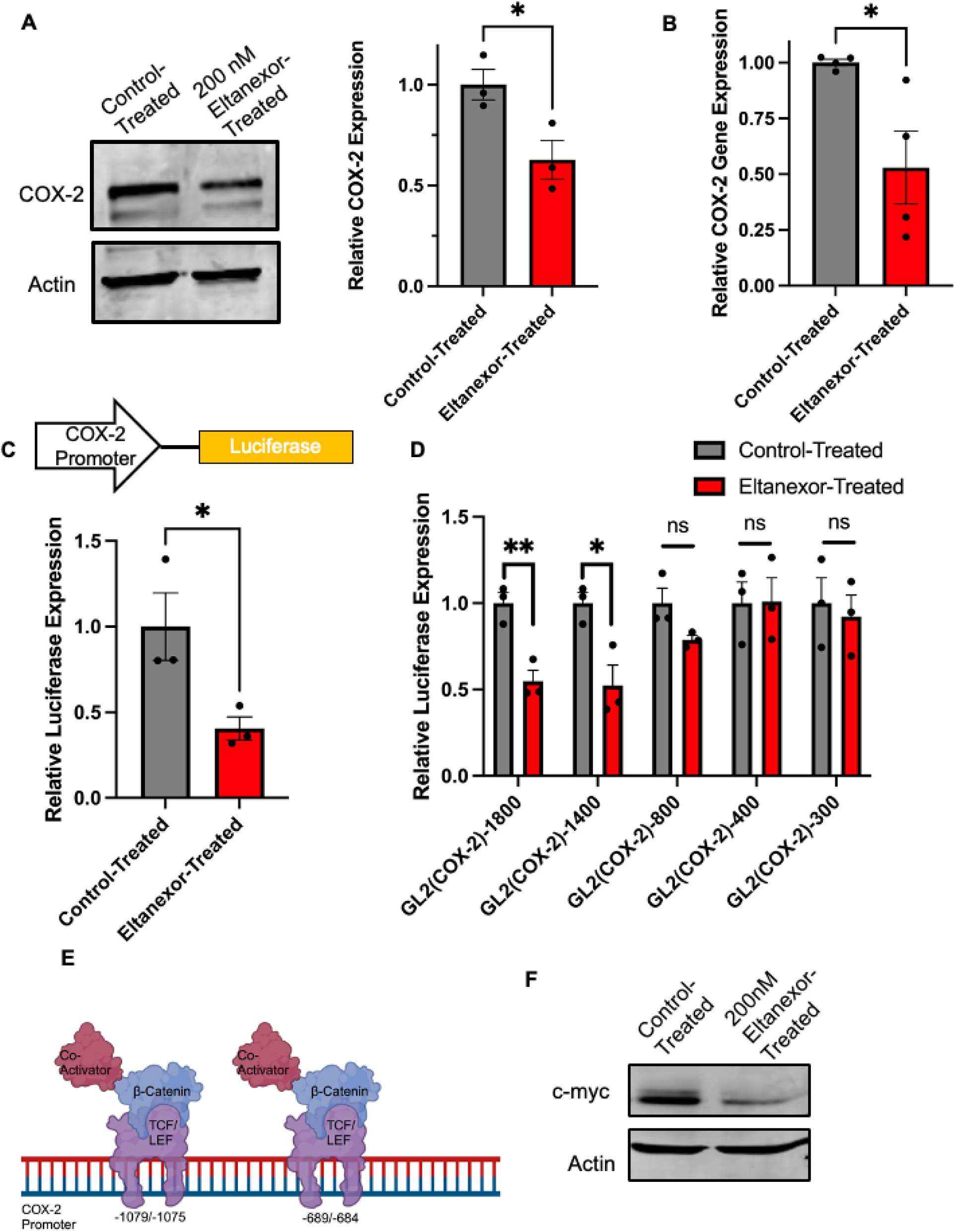
Eltanexor reduces COX-2 Expression. **(A)** Western blot probing for COX-2 protein in HCA-7 cells treated with either control or 200 nM Eltanexor for 48 hours. Actin was used as the loading control. The graph depicts the quantified densitometry of the western blot bands. The values have been normalized to the control-treated cells. The graph shows the mean of 3 independent experiments ± SEM. **(B)** COX-2 mRNA expression by qPCR in HCA-7 cells treated with DMSO control or 200 nM Eltanexor for 48 hours. The values are the mean of 4 experiments ± SEM. **(C)** HCT116 cells transfected with a plasmid containing the COX-2 promoter + Luciferase gene. The cells were subsequently treated with 200nM Eltanexor for 48 hours. The graph shows the mean Eltanexor-treated luciferase activity/ μg of protein normalized to DMSO-treated values of 3 independent experiments ± SEM. **(D)** HCT116 cells were transfected with COX-2 promoter + Luciferase Gene. The promoter region was deleted from full-length down to 300 base pairs. Cells were subsequently treated with DMSO or 200nM Eltanexor for 24 hours. The graph shows the mean luciferase activity/ μg of protein of Eltanexor-treated cells normalized to DMSO-treated values of 3 independent experiments ± SEM. **(E)** Schematic representation showing where Wnt/β-catenin signaling promotes COX-2 expression. **(F)** Western blot probing for c-myc in HCT116 cells treated with either control or 200 nM Eltanexor for 48 hours. *(*, p* ≤ *0.05; **, p* ≤ *0.01)*

To determine if Eltanexor can induce changes at the COX-2 promoter to regulate the protein’s expression, we utilized a luciferase plasmid containing the full-length COX-2 promoter (pGL3-COX2(−1840)). HCT116 cells were transfected with the pGL3-COX2(− 1840) reporter and then treated with 200nM Eltanexor for 48 hours. As shown in Figure 3C, Eltanexor can regulate COX-2 expression by impairing transcriptional activity at the promoter. To identify which key transcription factors Eltanexor treatment is altering, we performed a series of luciferase assays utilizing different regions of the COX-2 promoter. Eltanexor treatment significantly reduced luciferase activity by about 50% when regions −1800 (full length) and −1400 were transfected. When the −800 region was transfected, only about a 25% reduction in luciferase activity was observed. Furthermore, when the −400 and −300bp promoter regions were transfected, Eltanexor did not significantly reduce luciferase activity (**Figure 3D**). Interestingly, COX-2 has two functional T-cell factor/lymphoid enhancer factor (TCF/LEF)-response elements (TBE) sites. One of those TBE sites is at the −1079/−1074 region, and the other is at the −689/− 684 region of the COX-2 promoter [23–25]. TCF/LEF transcription factors interact with a crucial protein in CRC tumorigenesis, β-catenin, to execute Wnt signaling[25]. Our data suggests that Eltanexor may regulate COX-2 by impairing transcriptional activity at those TBE sites (**Figure 3E**). Additionally, we tested whether Eltanexor can regulate the expression of another gene controlled by β-catenin/TCF transcription, c-myc [26]. Interestingly, we show that Eltanexor can reduce c-myc protein expression (**Figure 3F**).

### Wnt/ **β**-catenin signaling is impaired with Eltanexor treatment

Along with being a commonly mutated pathway in sporadic CRC, the lack of regulation in the Wnt/ β-catenin signaling pathway results in the development of hundreds of polyps for people with FAP[27]. To determine if Eltanexor treatment can reduce Wnt/ β-catenin signaling, CRC cells possessing various Wnt/ β-catenin mutations were transfected with the Wnt/ β-catenin signaling luciferase reporter-TOPFlash. RKO possess wild-type Wnt/ β-catenin signaling pathway, DLD1 cells have a truncated APC protein at amino acid 1452 with WT β-catenin, while HCT116 cells have a β-catenin mutation due to ser45 that stabilizes the protein with WT APC[21]. The CRC cells were transfected with TOPFlash and then treated with Eltanexor for 24 hours. Our results revealed regardless of Wnt/ β-catenin mutational status, Eltanexor can reduce Wnt/β-catenin signaling (**Figure 4 A-C**). This data further suggests that XPO1 inhibition through Eltanexor can regulate COX-2 expression by impairing the Wnt/ β-catenin signaling pathway.

**Figure 4.**
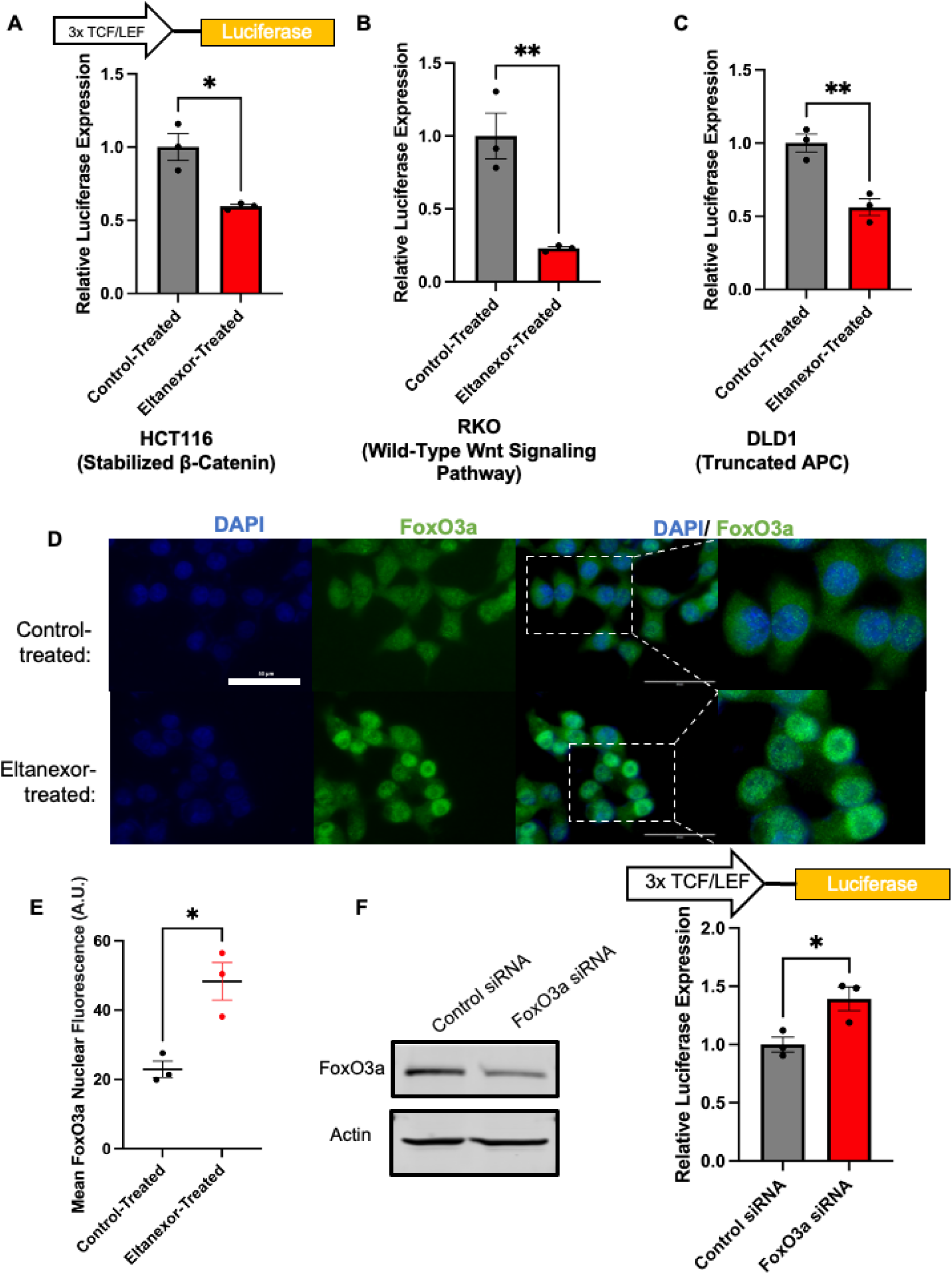
Eltanexor increases FoxO3a nuclear localization to modulate Wnt/β-catenin signaling. **(A-C)** HCT116, RKO, and DLD1 cells (all harboring different Wnt signaling phenotypes) were transfected with a TOPFlash reporter plasmid. Cells were subsequently treated with DMSO or Eltanexor. RKO and HCT116 cells were treated with 200nM Eltanexor, and DLD1 cells were treated with 400nM Eltanexor for 24 hours. Luciferase activity/ μg of protein was normalized to DMSO-treated luciferase expression for each respective cell line ± SEM. **(D)** HCT116 cells were treated for 48 hours with either 200 nM Eltanexor or DMSO. Subsequently, the cells were subject to immunofluorescent staining for FoxO3a (green). The nucleus was stained with DAPI (Blue). **(E)** The mean FoxO3a nuclear fluorescence for cells treated with DMSO (control) or 200 nM Eltanexor for 48 hours. The graph depicts the mean of 3 independent experiments ± SEM. At least 40 cells were analyzed for each independent experiment. The scale bars represent 50μm **(F)** In HCT116 cells, FoxO3a was knocked down by siRNA. Post-knockdown, the cells were transfected with a TOPflash reporter plasmid. The FoxO3a KD and TOPflash transfected cells were subject to immunoblotting to confirm FoxO3a KD. Actin was used as the loading control. The luciferase/ μg of protein value was normalized to control siRNA-treated cells ± SEM. **(G)** In HCT116 cells, FoxO3a was knocked down by siRNA. Then, the cells were treated with control or Eltanexor for 48 hours. Immunoblotting was used to determine changes in c-myc protein expression. *(*, p* ≤ *0.05; **, p* ≤ *0.01)*

### Eltanexor increases FoxO3a nuclear location which can modulate Wnt/ **β**-catenin signaling

To understand how XPO1 inhibition reduces Wnt/ β-catenin signaling, we first confirmed that Eltanexor does not alter β-catenin expression or intracellular localization in vitro (**Figure S5 A and B**). We next searched for proteins known to interact with XPO1 and β-catenin. In doing so, we identified the tumor suppressor, Forkhead Transcription Factor O Subfamily Member 3a (FoxO3a). Previous studies have shown that FoxO3a can modulate Wnt/β-catenin signaling in numerous cancer types[28–30]. Furthermore, in multiple cancer types, XPO1 inhibition increases nuclear FoxO3a intracellular localization both in vivo and in vitro[31,32]. In Figure 4 D and E, we show that in CRC cells treated with 200nM Eltanexor for 48 hours, FoxO3a increases in nuclear localization.

Next, we examined if FoxO3a can modulate Wnt/ β-catenin in CRC. When FoxO3a was knocked down (KD) in HCT116, there was approximately a 50% increase in Wnt/ β-catenin signaling by the TOPFlash Luciferase reporter (**Figure 4F**). Together, this data further highlights the ability of Eltanexor to contain FoxO3a in the nucleus to modulate Wnt/ β-catenin signaling.

### Apc*^min/+^* tumoroids are more sensitive to Eltanexor treatment compared to WT organoids

Apc*^min/+^* mice are a mouse model for FAP that harbor a nonsense mutation of the *APC* gene in the C57BL/6 background. The nonsense mutation results in a truncated APC protein[33–35]. Loss of heterozygosity of the heterozygous mutant leads to the development of tumors in the small intestine and colon of the mice by approximately four weeks of age[36].

To examine if Apc*^min/+^*mouse tumors are more sensitive to Eltanexor treatment compared to wild-type (WT) tissue, small intestinal tumoroids from Apc*^min/+^* mice and organoids from WT mice were created (**Figure 5A**).

**Figure 5.**
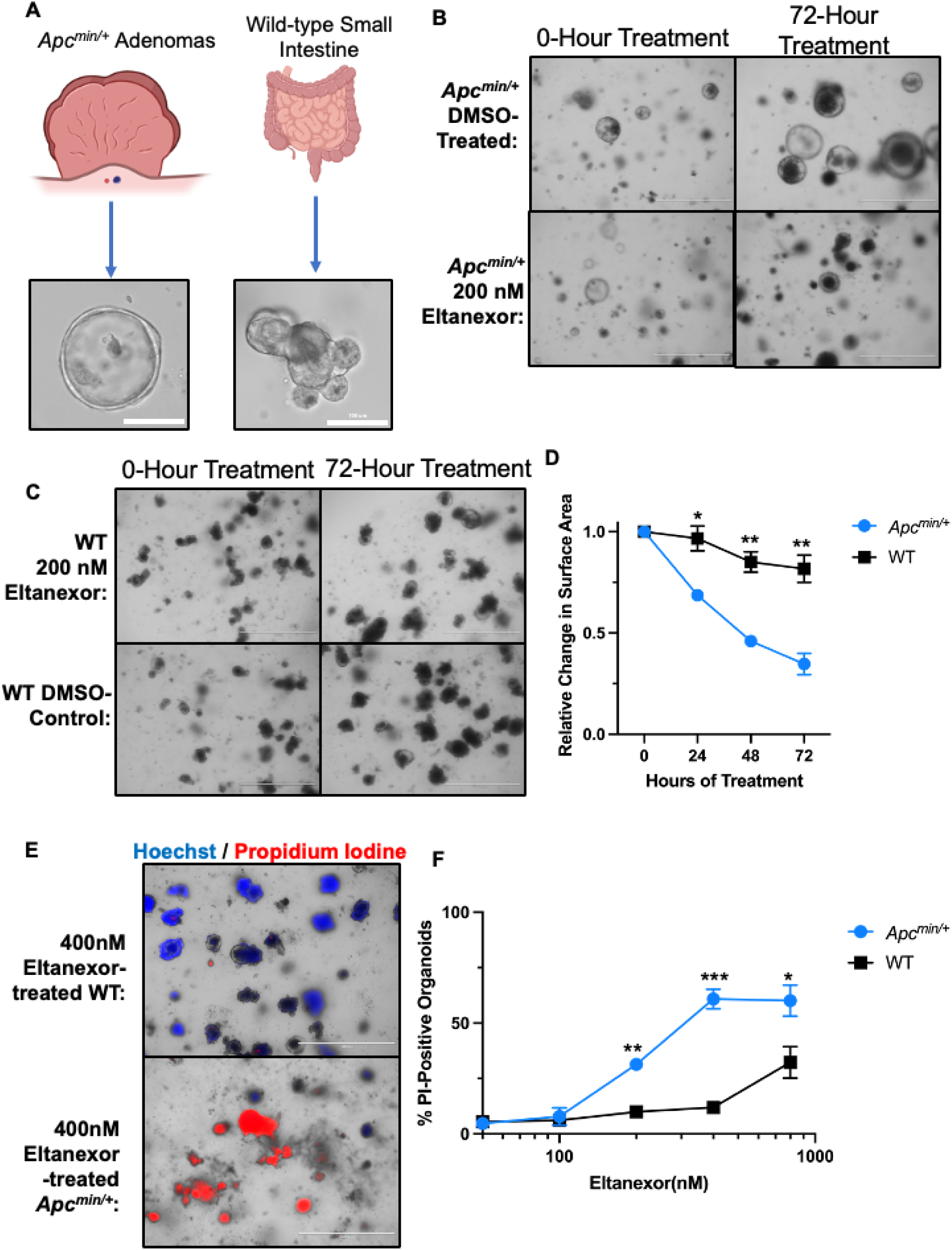
Eltanexor shows increased sensitivity to tumor-derived organoids compared to wild-type mouse small intestine organoids. **(A)** Representative photos of *Apc^min/+^*tumor-derived (tumoroids) and wild-type mouse small intestinal tissue-derived organoids. **(B)** Representative photos of tumoroids treated with 200nM Eltanexor for 72 hours. **(C)** Representative photos of wild-type organoids treated with 200nM Eltanexor for 72 hours. **(D)** Tumoroids and wild-type organoids were treated with a dosage of 200nM Eltanexor over 72 hours. Every 24 hours, the relative change in the surface area of each population was determined in ImageJ. The relative change is normalized to each population’s average surface area at hour 0. Values represent the mean of 3 individual experiments ± SEM. At least 50 organoids were measured at each time point under each treatment condition. **(E)** Representative photos of both organoid populations that were treated with 400nM Eltanexor for 72 hours. After 72 hours, each population was stained with 10 μg/mL of Hoechst and 10 μg/mL of propidium iodine. The organoids experiencing cell death will stain with propidium iodine. **(F)** Tumoroids and wild-type organoids were treated with DMSO, 50nM, 100nM, 200nM, 400nM, and 800nM Eltanexor for 72 hours. At 72 hours, the organoids were stained with 10 μg/mL of Hoechst and 10 μg/mL of propidium iodine. The graph represents the percentage of organoids from each population that were stained with propidium iodine under the various Eltanexor dosages. The values represent the mean of 3 individual experiments ± SEM. *(*, p* ≤ *0.05; **, p* ≤ *0.01; ***, p* ≤ *0.001)*

After five days, the tumoroids and organoids were treated with varying doses of Eltanexor. Images of both organoid populations were obtained every 24 hours for 72 hours. The images were analyzed to calculate the average size of organoids from each population and dosage. Our analysis has revealed that Eltanexor had a significant impact on the growth rate of tumoroids when compared to the effect it had on WT organoids (**Figure 5 B-D**). Furthermore, after 72 hours of treatment, the tumoroids and organoids were stained with Hoechst, which stains dead and alive organoids, and PI, which will only stain organoids undergoing cell death[37,38]. Then, we analyzed and scored the percentage of PI-positive organoids in each population, to determine percent organoid death. Our results reveal that the tumoroids experience a higher amount of organoid death at varying doses when compared to WT organoids (**Figure 5 E and F**). Altogether, these results indicate that the Apc*^min/+^*-derived tumoroids are more sensitive to Eltanexor treatment when compared to WT organoids. These results provide evidence that FAP model mice tumors are reliant upon XPO1 for survival and growth.

### Eltanexor acts as an effective chemopreventive agent in Apc*^min/+^* mice

XPO1 is consistently overexpressed in Apc*^min/+^* tumors (**Figure 6 A, S6 A**). To investigate the chemopreventive potential of Eltanexor, Apc*^min/+^* mice were treated with 10mg/kg of Eltanexor 3x/ week starting at the age of 4-5 weeks and for six weeks.

**Figure 6.**
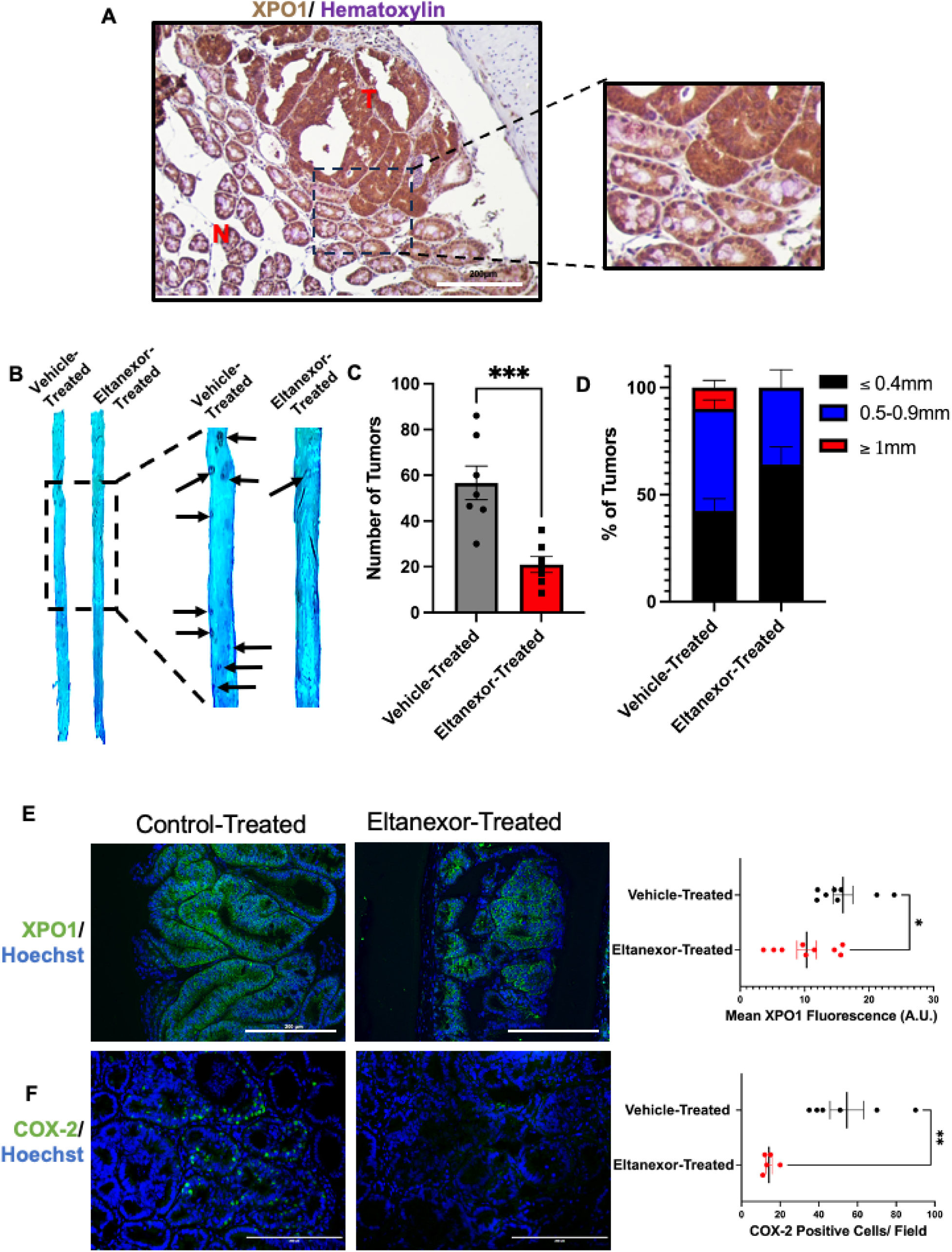
Eltanexor-treatment reduces tumor burden and size in Apc*^min/+^* mice. **(A)** Representative photo of *Apc^min/+^* mouse adenoma and adjacent normal colon IHC stained for XPO1. Nuclei are counterstained with Hematoxylin. The “T” label on the tissue represents tumor tissue, and the “N” label represents normal tissue. The scale bars represent 200μm. **(B)** Representative small intestine regions from Apc*^min/+^*mice that received vehicle or Eltanexor treatment. The tissue was fixed with 10% formalin and stained with 0.1% methylene blue. The arrows point to tumors present on the tissue. **(C)** Total intestinal tumor burden of Apc*^min/+^* mice that were treated from the age of 4 weeks to 11 weeks with 10mg***/***kg of Eltanexor or vehicle 3x/ week. The value represents the mean total intestinal tumor burden ± SEM of 7 different mice. **(E)** Frequency of <0.4mm, 0.5-0.9mm, and >1mm tumors in the vehicle-treated and Eltanexor-treated groups. **(E)** Representative images and quantifications of vehicle-treated and Eltanexor-treated tumors immunofluorescent stained for XPO1. Hoechst was used to stain the nucleus, and green represents the protein of interest. The quantifications are the mean fluorescent signal produced by an individual tumor. Three individual mice that received vehicle or Eltanexor treatment were used in the quantification. The values represent the mean XPO1 fluorescence ± SEM. The scale bars represent 200μm. **(F)** Representative images and quantifications of vehicle-treated and Eltanexor-treated tumors immunofluorescent stained for COX-2. Hoechst was used to stain the nucleus, and the green represents COX-2. The graph depicts the number of COX-2 positive cells per image. Three mice, vehicle-treated or Eltanexor-treated, were used in the quantification. The scale bars represent 200μm. *(*, p* ≤ *0.05; **, p* ≤ *0.01; ***, p* ≤ *0.001)*

Upon completion of the study, we examined the intestinal tract of the mice for changes in tumor burden and size. Our results reveal that mice that received Eltanexor drug treatment experienced an approximately 3-fold reduction in tumor burden (**Figure 6 B and C**). Effective chemopreventive compounds in Apc*^min/+^*can also reduce the weight of their abnormally large spleens[39]. In the Eltanexor-treated mice, the weights of the spleens were significantly smaller than the vehicle-treated mice and more similar in size to WT C67BL/6 mice (**Figure S6 B**). Additionally, Eltanexor-treated mice have a lesser frequency of tumors greater than 0.5mm in size (**Figure 6 D**). Our analysis shows that the Eltanexor-treated mice maintained a comparable body weight and intestinal tract to the vehicle-treated mice (**Figure S6 C and D**).

Furthermore, congruent with the effects we observed in vitro, the tumors of the mice treated with Eltanexor have a reduction in XPO1 (**Figure 6 E**). In the Eltanexor-treated tumors, we observe a reduction in the proliferation marker Ki67 **(Figure S6 E**). Finally, we also observe a consistent decrease of COX-2 in the Eltanexor-treated tumors (**Figure 6 F**). This observation further suggests the potential of Eltanexor to modulate Wnt/ β-catenin signaling in a mouse model to prevent CRC tumorigenesis.

## DISCUSSION

In this study, we demonstrate the significant impact XPO1 inhibition has on CRC tumorigenesis. Our findings reveal that Eltanexor treatment markedly reduces CRC cell viability by blocking XPO1’s interaction with cargo proteins and promoting subsequent XPO1 protein degradation. Furthermore, Eltanexor inhibition reduces COX-2 expression, a protein previously shown to be overexpressed in both CRC adenomas and adenocarcinoma [22]. COX-2 has a defined role in CRC tumorigenesis and is the predominant chemoprevention target in CRC[35,40,41]. Our results indicate that the Eltanexor-induced COX-2 reduction is mediated through modulation of the Wnt/ β-catenin pathway-one of the most mutated pathways in CRC [42].

Mutations in the *APC* gene result in aberrant Wnt/ β-catenin signaling in FAP patients, making the pathway the driver of tumorigenesis in these patients[43–45]. Our results show that XPO1 inhibition not only impairs numerous hallmarks of cancer but can also impair the signaling pathway that is the driver of tumorigenesis in FAP. Tumor-derived intestinal 3D cultures from the FAP mouse model, Apc*^min/+^*, showed high dependency on XPO1 for survival and growth, whereas low doses of Eltanexor had marginal effects on wild-type 3D cultures. Our preclinical evidence, derived from the treatment of Apc*^min/+^* mice, suggests Eltanexor could be an effective chemoprevention compound in FAP patients.

Previous studies and TCGA data suggest that XPO1 is consistently overexpressed in CRC[7]. Our analysis of XPO1 expression in multiple CRC stages and expression in the FAP model Apc*^min/+^*mouse adenomas suggest XPO1 overexpression is an early occurring event in CRC tumorigenesis. XPO1 is responsible for transporting 1000+ proteins from the nucleus to the cytoplasm[8]. Due to XPO1 interacting with many proteins, the inhibition of XPO1 can lead to the impairment of multiple different hallmarks of cancer[12–16], we hypothesized that XPO1 is a viable CRC chemoprevention target.

To target XPO1, SINE compounds have been developed and designed to create a slowly reversible covalent bond in XPO1’s cargo-binding groove that recognizes the protein’s specific nuclear export signal[13,15,46]. Eltanexor treatment reduced cell viability in multiple CRC cell lines with different genetic backgrounds within the nanomolar range. We additionally observed that Eltanexor treatment causes a dose-dependent reduction of XPO1. Furthermore, we confirmed these in vitro results with an in vivo xenograft study. Previous studies in other cancer cell types have determined this SINE compound-dependent XPO1 reduction is caused by the compound causing a conformational change in XPO1 that targets the protein for proteasomal degradation[32].

Correlations between XPO1 and COX-2 expression have been noted in ovarian cancer, where XPO1 inhibition reduces COX-2 expression[47]. Similar findings have been reported in CRC cells[48]. The COX enzymes contribute to the synthesis of prostaglandins (PG). The COX enzyme isoform, COX-2, is the rate-limiting step in prostaglandin E_2_ (PGE_2_) synthesis[35]. PGE_2_ can go on to perform signaling to promote tumorigenesis[49]. Previous studies have found that COX-2 inhibition can reduce tumor burden in Apc*^min/+^* mice and aids in reducing the risk for CRC in humans [50–53]. To examine Eltanexor as a chemopreventive compound, we tested whether the compound regulates COX-2 expression. Our study is the first to show that SINE compounds regulate COX-2 expression by altering the mRNA transcription of COX-2. These results provide more rationale suggesting Eltanexor will be an effective chemoprevention compound through its ability to regulate the protein that has been the chemoprevention standard for decades[41].

Eltanexor can reduce the transcriptional activity of the Wnt/ β-catenin signaling pathway, making the inhibitor an even more intriguing CRC chemopreventative compound. FoxO3a, which modulates the canonical Wnt/β-catenin signaling pathway by competitively binding to β-catenin in the nucleus[28–30], shows increased nuclear localization in Eltanexor-treated cells. This suggests the changes in FoxO3a subcellular localization contribute to the reduction of Wnt/ β-catenin signaling.

Interestingly, Apc*^min/+^* tumoroids exhibit increased sensitivity to Eltanexor treatment compared to WT organoids. The heightened sensitivity to Eltanexor suggests that Eltanexor may be well-tolerated and effective at limiting CRC tumorigenesis in an in vivo model. When we tested Eltanexor in the Apc*^min/+^* mice, we were encouraged to see similar results as our ex vivo study. The Apc*^min/+^* mice given Eltanexor experienced reduces tumor burden and size while being well-tolerated. These results show that Apc*^min/+^* tumors rely on XPO1 for viability and growth.

This study provides compelling preclinical evidence for Eltanexor’s effectiveness as a chemoprevention compound for FAP patients. Future studies should aim to further evaluate the safety and efficacy of Eltanexor as a chemopreventive agent in FAP patients. Additionally, diseases such as Inflammatory Bowel Disease and Lynch Syndrome, which increase the risk of CRC, also warrant the exploration of novel chemoprevention compounds like Eltanexor [54,55]. Given XPO1’s invovlvement in numerous pathways related to CRC tumorigenesis, Eltanexor holds promise as a broad-spectrum chemopreventive agent in high-risk patient populations.

## Supplemental Figures

**Figure S1.**
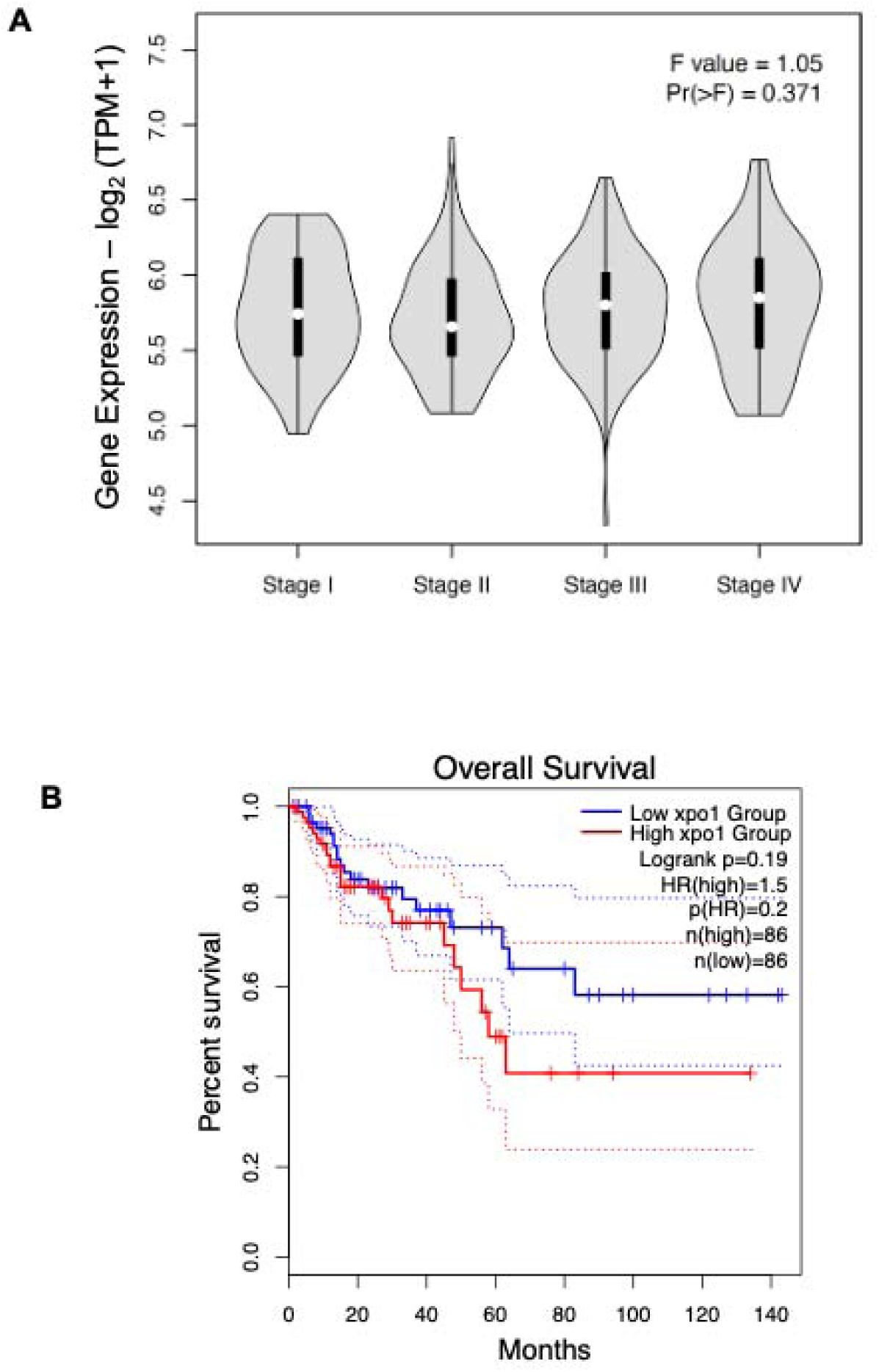
XPO1 expression is similar amongst all stages of CRC and overexpression trends towards worst prognosis in MSS CRC. **(A)** Analysis of XPO1 expression amongst CRC tumor stages derived from GEPIA (http://gepia2.cancer-pku.cn/#analysis). **(B)** Kaplan-Meier plot derived from GEPIA showing colon adenocarcinoma (COAD) microsatellite stable patient survival with high or low XPO1 expression (GEPIA: http://gepia2.cancer-pku.cn/#survival)

**Figure S2.**
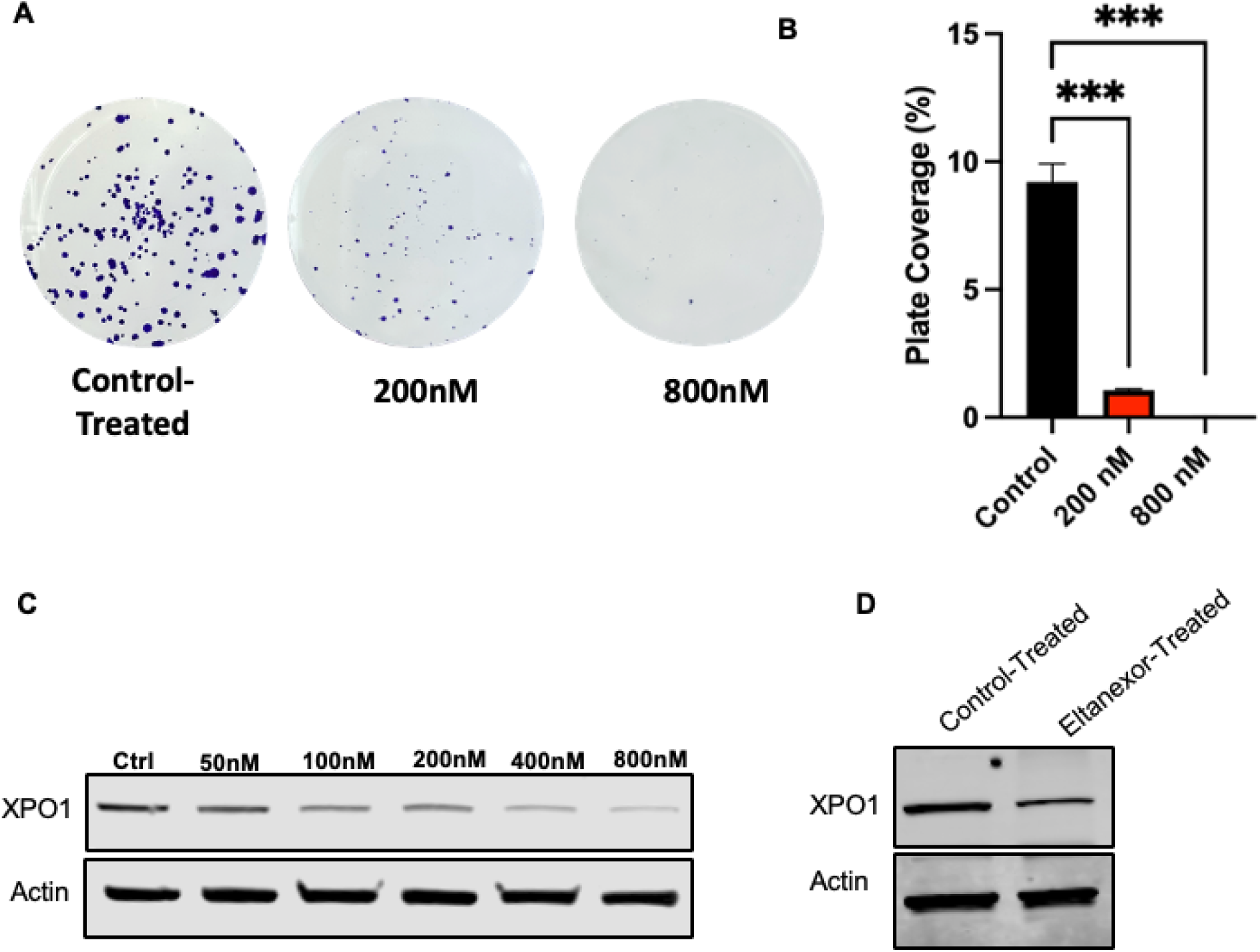
(A) Eltanexor treatment reduces XPO1 protein expression in multiple CRC cell lines and prevents colony formation. (A, **B)** HCT116 cells were plated with 500 cells/ well and treated with control, 200nM Eltanexor, or 800nM Eltanexor for 72 hours. The cells were cultured for an additional 8 days and stained with 0.1% crystal violet solution. The mean percent of the plate covered by colonies of 3 independent experiments ± SEM was determined. **(C)** XPO1 expression in HCT116 cells treated with varying doses of Eltanexor for 48 hours. **(D)** XPO1 expression in HCA7 cells treated with 200nM Eltanexor for 48 hours. *(***, p* ≤ *0.001)*

**Figure S3.**
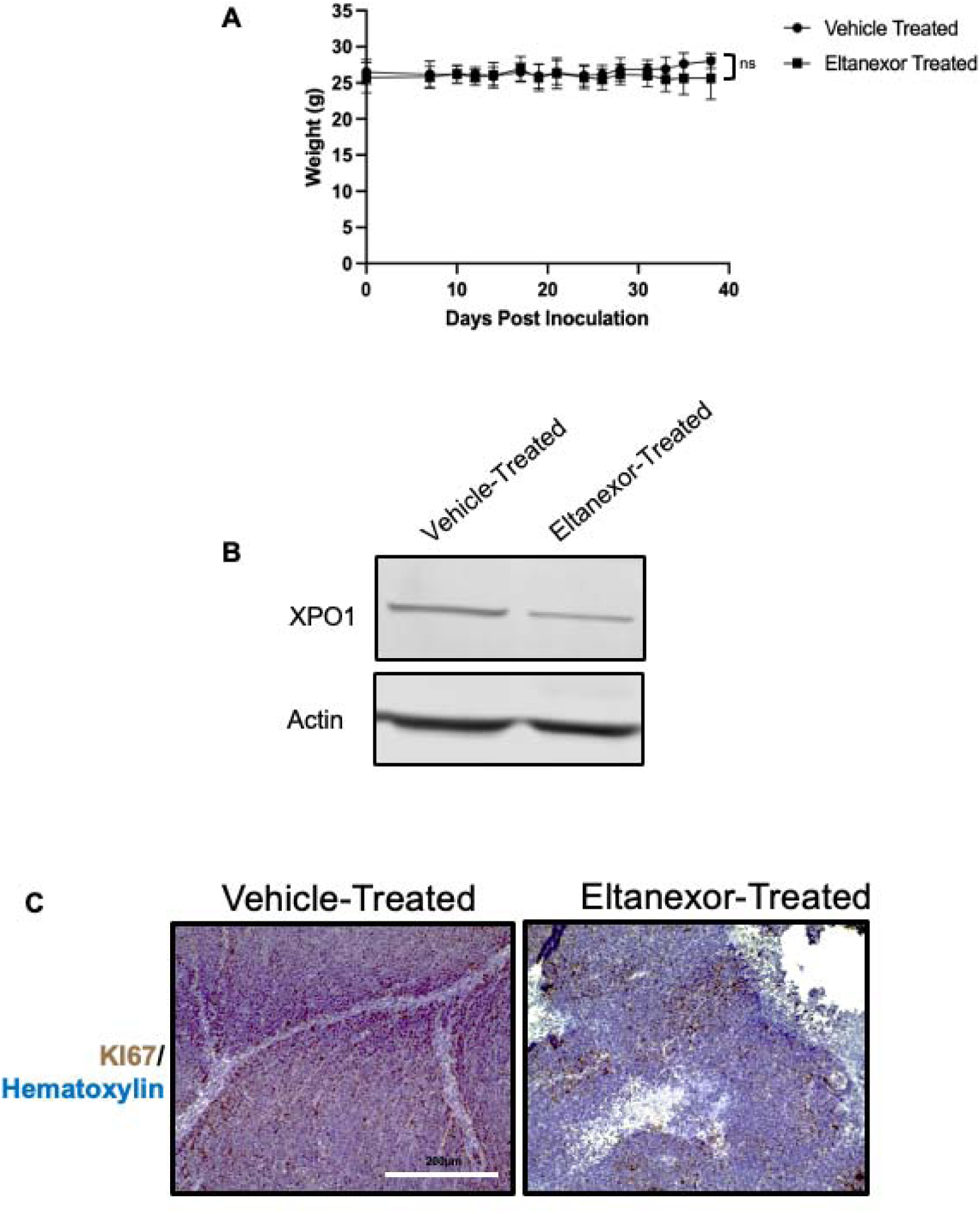
Eltanexor treatment reduces HCT116 xenograft Growth. **(A)** Athymic mice injected with HCT116 cells and treated with either vehicle or Eltanexor weights over the course of the study. Each data point represents the mean mouse weight ± SEM. **(B)** Representative XPO1 protein expression from tumors derived from HCT116 xenograft mice treated with either vehicle or Eltanexor. **(C)** IHC detection of Ki67 in HCT116 xenograft tumors. Representative sections were stained for Ki67 (brown) and counterstained with hematoxylin (blue). The scale bars represent 200μm.

**Figure S4.**
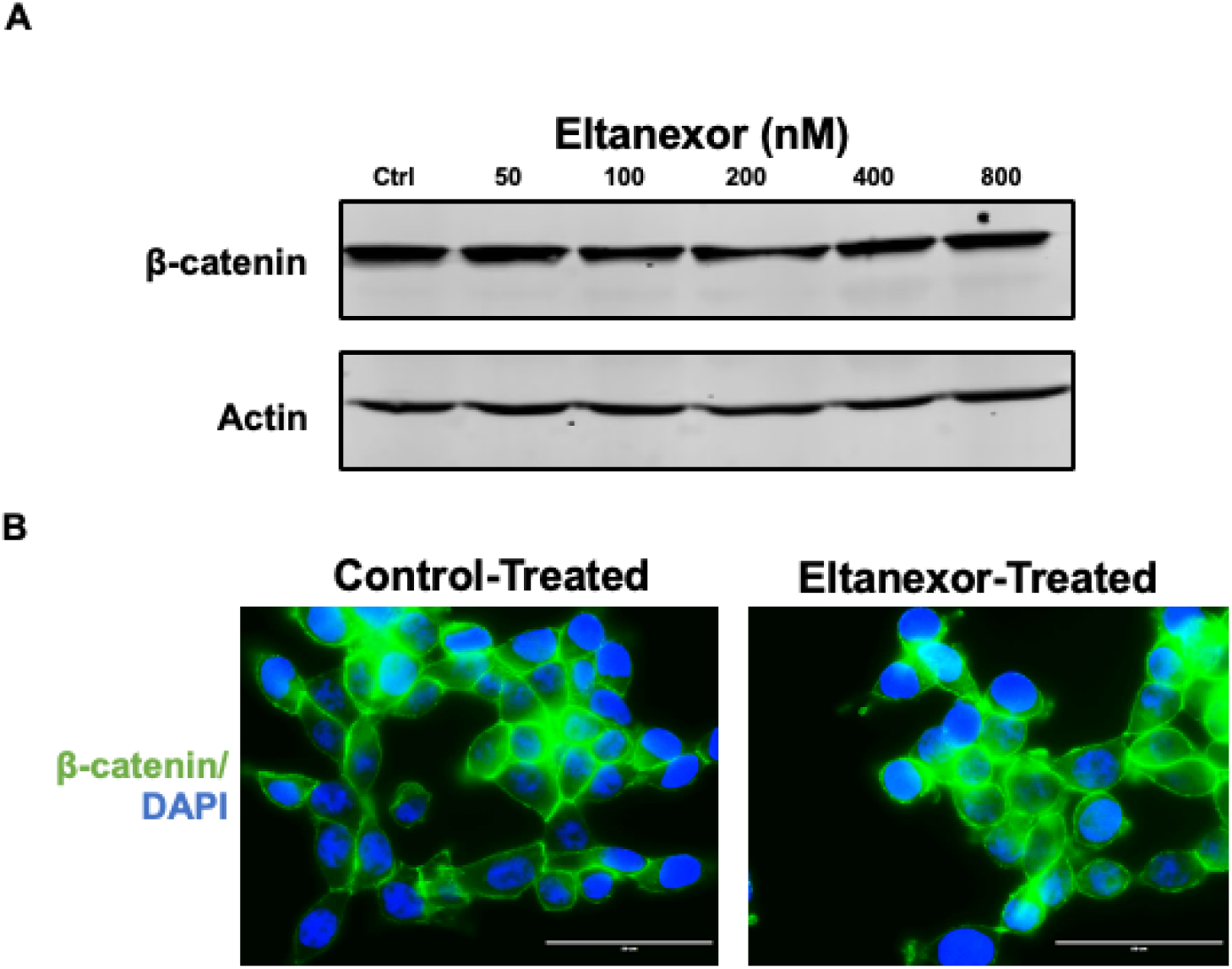
Eltanexor does not changes. β**-catenin protein expression or intracellular localization. (A)** HCT116 cells were treated with varying doses of Eltanexor or DMSO (control) for 24 hours. Subsequently, the cells were subjected to immunoblotting for changes in β-catenin protein expression. **(B)** HCT116 cells were treated with 200nM Eltanexor or DMSO (control) for 48 hours. The cells were subject to immunofluorescent staining for FoxO3a (green). The nucleus was stained with DAPI (Blue).

**Figure S5.**
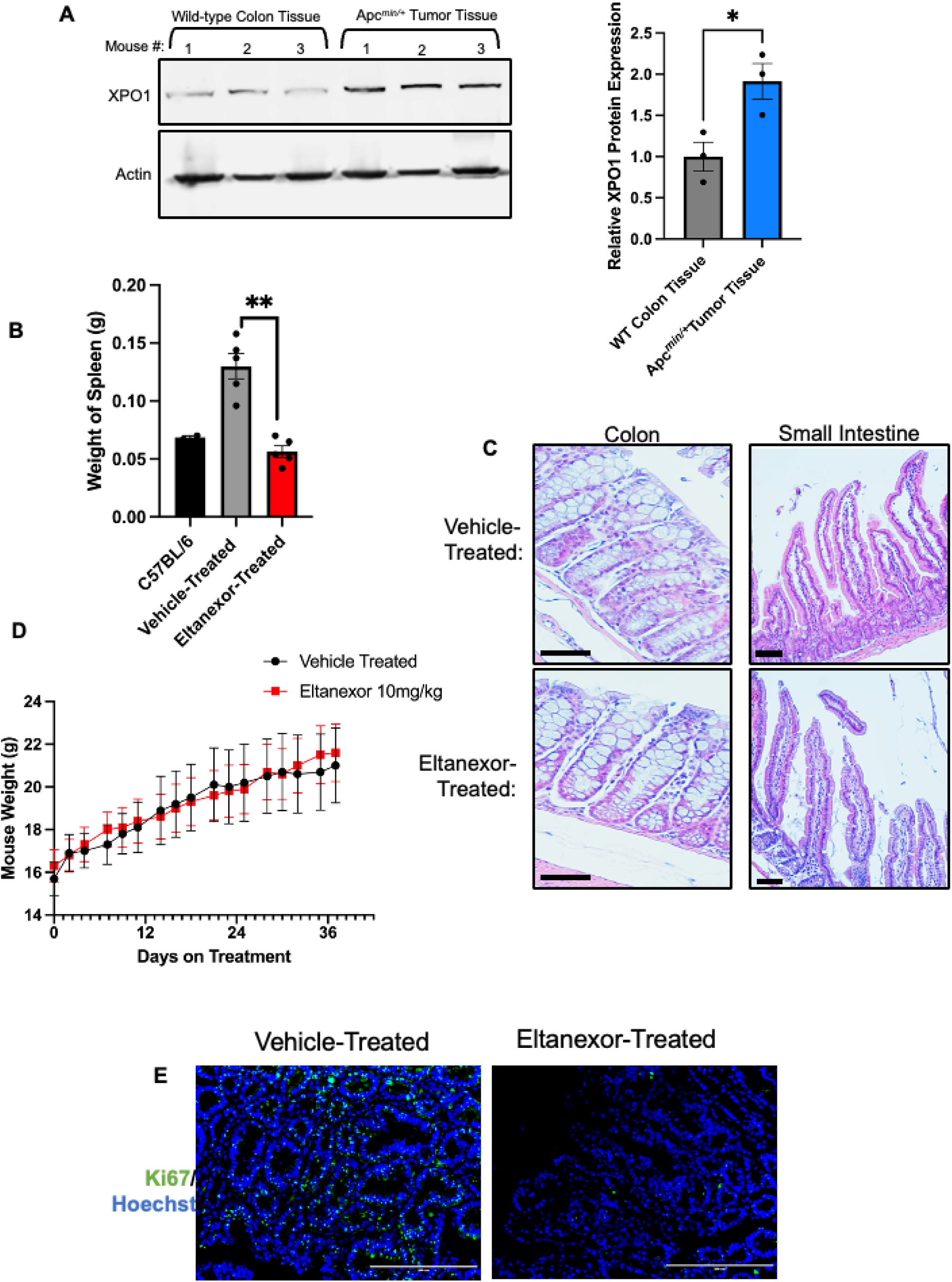
Eltanexor-treatment is well-tolerated in Apc*^min/+^* mice. **(A)** Colon epithelial tissue from three wild-type C57BL/6 mice was collected, while colon tumors were removed from three Apc*^min/+^* mice. The tissues were then ran on a western blot to evaluate changes in XPO1 protein, with actin as the loading control. The graph shows the relative protein expression of XPO1 normalized to WT mice. **(B)** Apc*^min/+^* mice were treated with either vehicle or 10 mg/kg of Eltanexor 3 days a week for 6 weeks. The weight of each mouse was recorded every Monday, Wednesday, and Friday. The values represent the mean weight of the mice from each group on the given treatment date ± SEM. **(C)** Post-treatment, intestinal tissue was formalin-fixed paraffin-embedded. Representative Hematoxylin & Eosin (H&E)-stained mid-colon and proximal small intestine from mice either treated with vehicle or Eltanexor. The hematoxylin (purple) represents the nucleus, while the eosin (pink) represents the cytoplasm. Scale bars represent 50μm. **(D)** The mean spleen weight of C57BL/6 mice, vehicle-treated Apc*^min/+^*mice, and Eltanexor-treated Apc*^min/+^* mice ± SEM. Spleen weights of the C57BL/6 mice were age-matched to the treated mice. **(E)** Formalin-fixed paraffin-embedded tissue from vehicle-treated and Eltanexor-treated was immunofluorescent stained for Ki67 (green), and the nuclei were stained with Hoechst (blue). Scale bars represent 200 μm. *(*, p* ≤ *0.05; **, p* ≤ *0.01; ***, p* ≤ *0.001)*

